# Rod photoreceptors control the ON vs OFF polarity of cone-signaling neurons

**DOI:** 10.1101/2025.06.26.661749

**Authors:** Deborah Langrill Beaudoin, Abdul Rhman Hassan, Angela Shehu, Jeremy M. Bohl, Yumiko Umino, Eduardo C. Solessio, Chase B. Hellmer, Tomomi Ichinose

**Affiliations:** Department of Ophthalmology, Visual and Anatomical Sciences, Wayne State University School of Medicine, Detroit, MI, USA; Department of Ophthalmology, State University of New York, Upstate Medical University, Syracuse, NY, USA

## Abstract

A fundamental feature of the visual system is its ability to detect image contrast. The contrast processing starts in the first synapse of the retina where parallel pathways are established to compute contrast to bright (ON pathway) and dark (OFF pathway) objects, separately transferred to morphologically identified ON and OFF cells throughout the visual system. Here, we found that response polarity in ON and OFF neurons is not fixed but rather switches dynamically to the opposite sign. The switch was not observed in rod-knockout mice, indicating that rods generate the polarity switch. We determined that neither horizontal cells nor rod-signaling pathways were responsible for the switch. Instead, we discovered that EAAT5 glutamate transporters located at photoreceptor terminals were required to produce the polarity switch. Our findings provide a new perspective on the adaptive properties of neural networks and their ability to encode contrast across the visual dynamic range.

## Main

The retina in the eye is the first component of the visual system, capturing and processing a wide variety of image signaling. Various image features, such as color and motion, are encoded into separate neural networks, consisting of numerous types of retinal neurons and synapses. The retinal excitatory pathway starts from the first-order neurons, the rod and cone photoreceptors. Signals are transferred to second-order neurons, bipolar cells, and then to third-order neurons, ganglion cells. As visual signals are transferred through the excitatory pathway, they are tuned by multiple antagonistic systems. Antagonistic center and surround mechanisms occur via horizontal and amacrine cells, which spatially tune excitatory signals. Additionally, rods and cones induce suppressive interactions at mesopic light conditions when both photoreceptors are active ^1–4^. Furthermore, ON and OFF signaling pathways separately convey two opposite signs of excitatory signals.

ON and OFF signaling pathways encode positive (bright) and negative (dark) contrasts, respectively. Approximately half of the inner retinal neurons, bipolar, amacrine, and ganglion cells, are categorized as ON cells, excited (or depolarized) by a light stimulus. In contrast, OFF cells are excited by a dark stimulus or shut off (or hyperpolarize) at the light onset. As the signals are transferred from second to third-order neurons, ON and OFF signaling transmission occurs in the different sublaminae of the inner plexiform layer (IPL) in parallel. This morphological and physiological relation was discovered originally by Famiglietti et al. a half-century ago ^5, 6^, and has been well agreed upon and supported by subsequent retinal studies^7–12^.

We previously recorded from ON and OFF cone bipolar cells using mouse retinal slice preparations. Using this technique, we rarely observed violations of the ON/OFF morphological-physiological relations among more than 100 bipolar cell recordings completed ^8, 9^. However, when we recently used wholemount retinal preparations, we unexpectedly observed that many second- and third-order neurons exhibited signal polarity switches. Interestingly, many of those neurons switched their signs as the ambient light level changed. In the present study, we pursued the mechanisms underlying the visual signaling polarity switch.

## Results

We conducted patch clamp recordings from starburst amacrine cells (SACs) in wholemount tissues from Ai9; ChAT-cre mice, in which SACs were labeled with tdTomato. We confirmed the SAC recording by injecting a green fluorescent dye, Alexa 488, through the intracellular pipette and morphological examination, which revealed several symmetric primary dendrites having wavy branches and numerous varicosities in the distal branches (Fig. 1a). Physiologically, the SAC recording was confirmed by transient potassium currents without prominent sodium currents in response to voltage steps (Fig. 1b), consistent with previous reports ^13^. We targeted both ON and OFF SACs whose somas reside in the ganglion cell layer and inner nuclear layer, respectively ^14^. We recorded light-evoked excitatory postsynaptic potentials (L-EPSPs) in ON SACs and found that a fraction of the cells depolarized at light onset, as is expected for ON SACs (n=24, Fig. 1ci). However, unexpectedly, we observed that many other ON SACs hyperpolarized at the light onset (n=34, Fig. 1cii). Such a hyperpolarizing response is typically observed in OFF cells, but their somas were located in the ganglion cell layer, consistent with an ON SAC type.

**Fig. 1.**
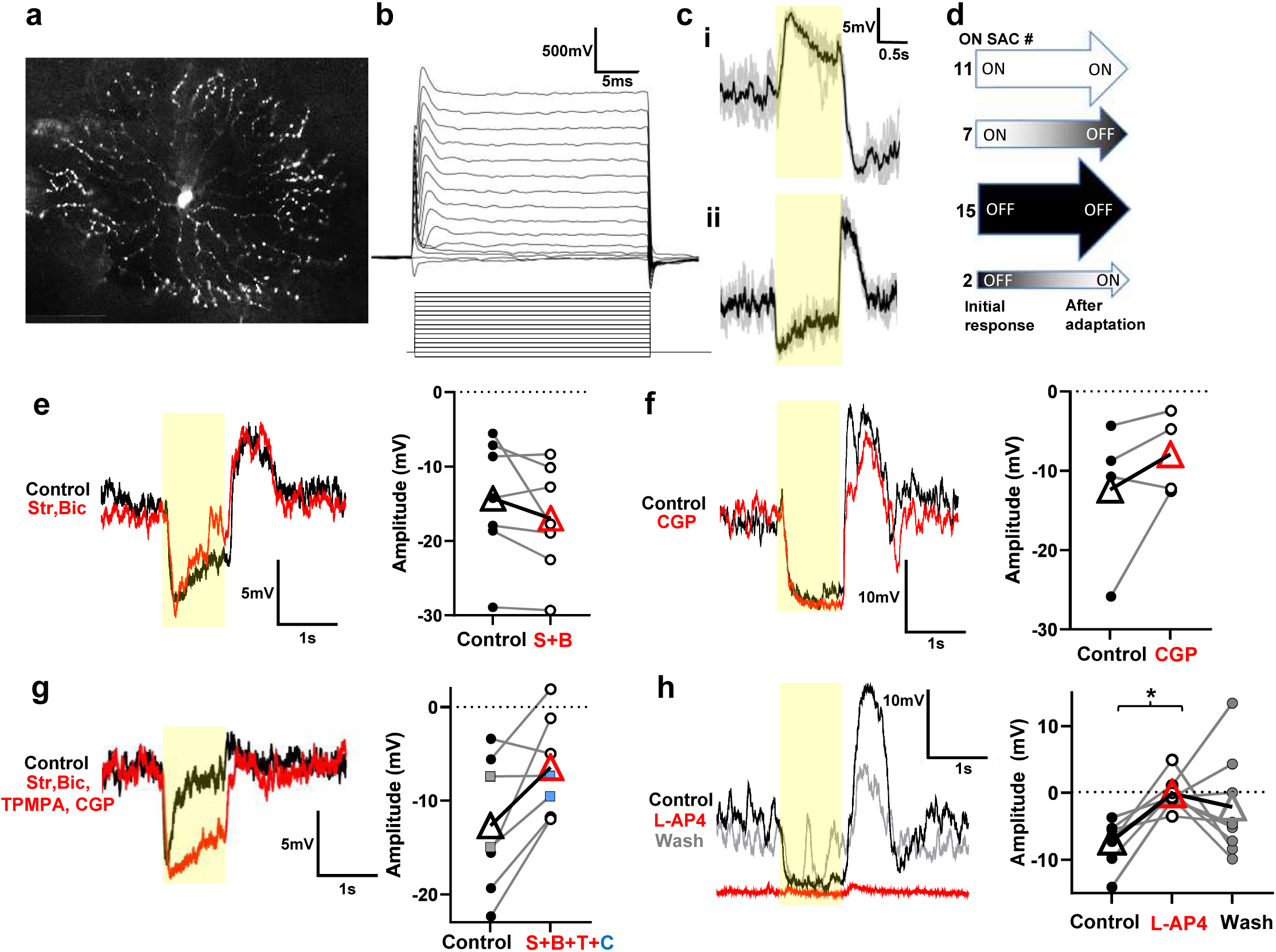
ON SACs exhibit polarity switch, which was not evoked by light adaptation, antagonistic surround, or OFF signaling crossover. **a.** An ON SAC labeled with an intracellular florescent dye, sulforhodamine b. **b.** A family of whole cell currents in response to 10 mV depolarizing steps from −70 to +20 mV, showing transient outward potassium currents without sodium currents. **c. i**. A representative depolarizing response to 1-second 20% contrast step light stimulus (yellow bar) at mesopic conditions. Individual traces are shown in gray, and the average response in black. **ii**. Atypical hyperpolarizing light-evoked response of another ON SAC to same stimulus as (**i**). **d.** Representation of ON SAC response polarities that exhibit initial depolarizing (white) or hyperpolarizing (black) responses with progression of responses after light adaptation. **e.** A hyperpolarizing L-EPSP in an ON SAC before (black) and after (red) application of 1 µM strychnine and 50 µM bicuculline to block glycine and GABA-A receptors. A graph summarizes L-EPSP amplitudes from 7 ON SACs, showing no effects on polarity switch. **f.** A hyperpolarizing ON SAC before (black) and after (red) application of 10 µM CGP-52432, a GABA-B antagonist. A graph summarizes L-EPSP amplitudes from 4 ON SACs. **g.** A hyperpolarizing ON SAC before and after application of 50 µM TPMPA, a GABA-C antagonist, in addition to strychnine, bicuculline, and CGP-52432. The white circles represent responses after application of Str, Bic & TPMPA, while the blue squares represent the latter 3 blockers + CGP. A graph summarizes L-EPSP amplitudes from 7 ON SACs. **h.** A hyperpolarizing ON SAC before and after application of 10 µM L-AP4, an mGluR6 agonist, to block mGluR6 signaling. A graph showing L-EPSP amplitudes for three conditions from 8 ON SACs. In all graphs for this and subsequent figures, circles indicate the amplitudes of individual cells and large triangle symbols indicate average of the responses. Paired t-tests were used for **e**, **f** & **g**, while Repeated ANOVA with Tukey’s multiple comparison was used for **h**. Asterisks represents statistically significant effects.

ON and OFF polarity switches in SACs were reported previously ^15^, where the authors observed that the light response polarity switch occurs after light adaptation during the recording. However, in our recordings, the polarity switch occurred randomly and was not driven by light adaptation in mesopic conditions (Fig. 1d): approximately half of ON SACs exhibited an ON response, and others showed an OFF response at the beginning of recordings. Some SACs switched from ON to OFF or OFF to ON, while others remained as either ON or OFF during the recordings. Hence, the previous findings ^15^ did not explain our results, and thus, we investigated the source of the ON and OFF polarity switches.

### The polarity switch was not induced by antagonistic surround, nor by OFF signaling crossover

We first examined whether the unexpected polarity changes were generated by the antagonistic surround mediated through amacrine cells. Amacrine cells release either GABA or glycine to inhibit neurons in the IPL, and ON SACs express both GABA-A and glycine receptors ^16–18^. We recorded from a hyperpolarizing ON SAC, and found that application of the glycine and GABA-A receptor blockers, strychnine (1 µM) and bicuculline (50 µM), increased the spontaneous activity, but did not change the polarity switch. The cell continued to hyperpolarize in response to light (Fig. 1e). SACs also express GABA-B receptors ^19^. However, when we applied a GABA-B receptor antagonist, CGP-52432 (10 µM), to an ON SAC that exhibited a polarity switch, we found that CGP slightly changed the EPSP amplitudes, but it did not affect the polarity switch (Fig. 1f). We also tested the effects of blocking GABA-C receptors using TPMPA (50 µM). GABA-C receptors are not expressed by SACs but by presynaptic bipolar cells ^20^. Application of a cocktail of GABA and glycine receptor antagonists (bicuculline, CGP52432, TPMPA, strychnine) induced spontaneous activity but had no effects on the polarity switch (Fig. 1g). Altogether, these results indicate that the antagonistic surround did not induce the ON SAC polarity switch.

We then examined whether an OFF signaling crossover induced the polarity switch in ON SACs. We used an mGluR6 agonist, L-AP4 (10 µM), which blocks synaptic transmission from photoreceptors to ON bipolar cells, blocking signaling exclusively in the ON pathway ^21^. The application of L-AP4 abolished the hyperpolarizing light response in ON SACs (Fig. 1h), indicating that the polarity switches in ON SACs are induced by ON pathway signaling.

The polarity switch we observed hyperpolarized an ON SAC to near −70 mV. We examined whether potassium channels played a role in the hyperpolarizing response because the equilibrium potential of potassium channels is near −86 mV based on the composition of our extracellular and intracellular solutions. SACs express Kv3 channels, a type of voltage-gated potassium channels, near the soma ^13^. We applied TEA (1 mM) in the bath solution to block the Kv3 channels, which generated massive spontaneous activity, but the polarity switch remained (Extended Data Fig. 1a). We also applied TEA (10 mM) in the recording pipette to block the channel intracellularly, but did not affect the polarity switch (Extended Data Fig. 1b). We furthermore tested a cesium intracellular solution to block a broad spectrum of potassium channels, and this also failed to remove the polarity switch (n=9 SACs, data not shown). These data indicate that voltage-gated potassium channels did not cause the polarity switch in ON SACs.

### Rod-cone interaction induced a polarity switch in ON SACs

Szikra, et al. ^22^ reported that rods receive cone input through horizontal cells in the mesopic to photopic conditions, inducing the polarity switch in a rod-signaling pathway. Therefore, we examined whether rod-cone interaction induced the polarity switch. L-EPSPs were recorded from ON SACs in response to a wide range of light intensities. To our surprise, ON-SACs exhibiting the polarity switch in mesopic conditions gradually and consistently corrected the sign as the light levels increased to photopic ranges (Fig. 2a, p<0.01). We also recorded L-EPSPs from tdTomato-expressing OFF SACs whose somas reside in the inner nuclear layer (INL). OFF SACs similarly exhibited a polarity switch: depolarizing at mesopic backgrounds but hyperpolarizing in photopic conditions (Fig. 2b, p<0.01). These results suggest that rod and cone interactions in the mesopic light conditions caused the polarity switch.

**Fig. 2.**
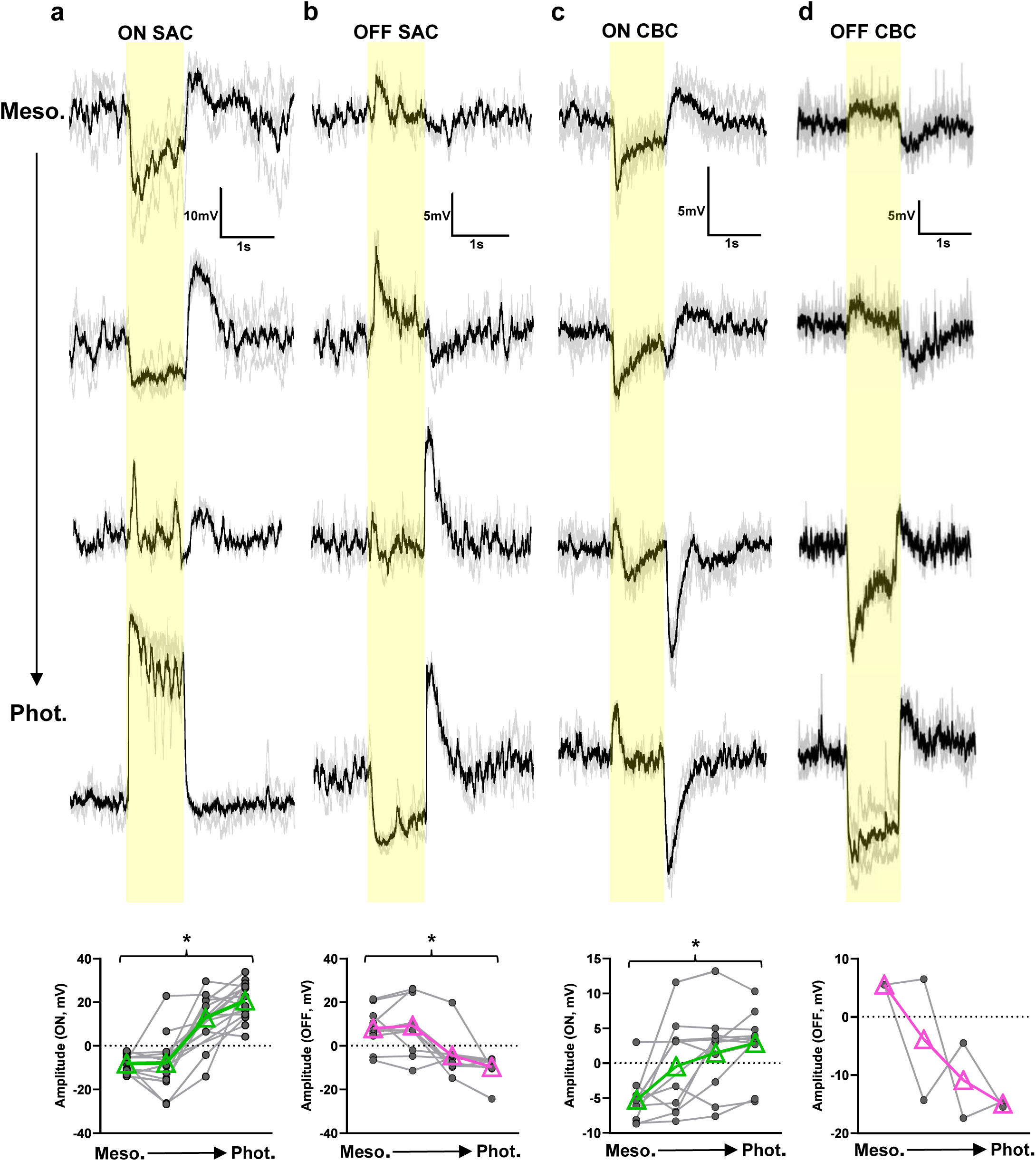
Increasing luminance levels corrected polarity switch in ON and OFF SACs and CBCs. **a.** An ON SAC exhibited hyperpolarized L-EPSPs at mesopic light levels (yellow bar indicates the timing of light stimulus) and became more depolarized as the background light levels increased to photopic range. In all panels, the black trace is the average of the gray trials. A graph at the bottom showing L-EPSP amplitudes in 11 ON SAC cells from mesopic to photopic luminance. Black circles indicate response amplitude in individual ON SAC, and green triangles show average responses. **b.** An OFF SAC depolarized at the light onset in the mesopic conditions and became more hyperpolarized as the background light levels increased to photopic. A graph at the bottom showing 8 OFF SACs exhibiting similar responses (black circles show individual OFF SACs, and magenta triangles show average responses). **c.** An ON CBC exhibited hyperpolarizing L-EPSP at mesopic levels, which became more depolarized as the background light levels increased to photopic. A graph at the bottom showing 9 ON CBCs with similar progression over increase in luminance levels. **d.** An OFF CBC exhibited depolarizing L-EPSP in mesopic conditions became more hyperpolarized as background light levels increased to photopic. A graph at the bottom shows 2 OFF bipolar cells with similar progression over increase in luminance levels. All statistical tests in this figure are One-Way ANOVA with Tukey’s multiple comparison. Asterisks represents statistically significant effects.

We tested whether horizontal cells mediated the rod-cone interactions. Horizontal cells’ output to cones involves pH change of the photoreceptor synapse, which can be suppressed by applying the pH buffer, HEPES, in the bath solution ^23^. HEPES slightly changed the amplitude of the light response; however, the polarity switch was not corrected (Extended Data Fig. 1c,d). Although HEPES did not affect the polarity switch in ON SACs, we examined horizontal cells’ contribution using different methods in later experiments.

### ON and OFF cone bipolar cells exhibited polarity switch

If rod-cone interaction induced the polarity switch in ON SACs, bipolar cells, which are upstream neurons for ON SACs, should also show a polarity switch. Our laboratory previously conducted patch clamp recordings from more than 100 ON and OFF cone bipolar cells (CBCs) in slice preparations (250 µm thick), in which ON CBCs, defined by the axon terminals ramifying in the IPL ON sub-laminae, depolarize to light onset, whereas morphologically OFF CBCs hyperpolarize to light onset ^8, 9^. We rarely observed violations of the ON and OFF signs and morphological types, consistent with the fundamental rule in the field ^5, 6^. Therefore, we were initially skeptical about ON and OFF polarity switches in CBCs.

Unlike our previous experiments working in retinal slice preparations, here we recorded from bipolar cells using wholemount retinal preparations, the same preparations we used for SAC recordings. We previously established this difficult approach for bipolar cell patch clamp and imaging in wholemount preparations ^24, 25^. In mesopic conditions, we observed that many ON CBCs hyperpolarized to light onset, exhibiting a polarity switch (Fig. 2c). Similar to ON SACs, light-evoked hyperpolarization in mesopic conditions was converted to depolarization as the light level increased to the photopic range (Fig. 2c). Similarly, OFF CBCs showed a polarity switch in mesopic conditions, which was changed to the correct sign in photopic conditions (Fig. 2d). Approximately half of ON and OFF CBCs exhibited polarity switch without clear type-dependency (Table 1, WT). We occasionally observed polarity switch occurring in an opposite way, correct sign in mesopic and switched in photopic conditions (Extended Data Fig. 2a).

**Table 1.** Summary of ON and OFF bipolar cell numbers exhibiting a polarity switch between mesopic and photopic light conditions for different genetic mouse models. WT = wildtype, RKO = rod knockout, CKO = cone knockout, UK type = unknown type.

|  | Bipolar type | Polarity switch |  | Non-switch |  |
| --- | --- | --- | --- | --- | --- |
| WT | ON | 13 | Type 5: (5)<br>Type 6/7: (8) | 7 | Type 7: (1)<br>Type 5: (1)<br>Type 6/7: (2)<br>UK Type: (3) |
|  | OFF | 11 | Type 3/4: (9)<br>UK Type: (2) | 15 | Type 1/2: (6)<br>Type 3/4: (7)<br>UK Type: (2) |
| RKO<br>(Gnat1<sup>-/-</sup>) | ON | 0 |  | 5 | Type 5: (3)<br>Type 6: (1)<br>UK Type: (1) |
|  | OFF | 0 |  | 5 | Type 1/2: 1<br>Type 3: 1<br>UK Type: 3 |
| CKO<br>(Cnga3<sup>-/-</sup>) | ON | 10 | Type 5: (2)<br>Type 6/7: (2)<br>UK Type: (6) | 3 | Type 5: (1)<br>Type 6/7: (1)<br>UK Type: (1) |
|  | OFF | 10 | Type 1/2: (1)<br>Type 3/4: (1)<br>UK Type: (8) | 3 | UK Type: (3) |
| CKO<br>(Gnat2<sup>-/-</sup>) | ON | 8 | Type 5: (4)<br>Type 6/7: (1)<br>UK Type: (3) | 0 |  |
|  | OFF | 10 | Type 1/2: (1)<br>Type 3/4: (3)<br>UK Type: (6) | 3 | Type 1/2: (1)<br>Type 3/4: (2) |

Interestingly, rod bipolar cells did not show a polarity switch but responded both in mesopic and photopic conditions (Extended Data Fig. 2b). Our previous and current data suggest that the polarity switch in bipolar cells was induced by a mechanism only in wholemount preparations, such as lateral circuitry. Because wholemount preparations are more physiological than the slice, we postulate that the polarity switch is a physiological event.

### Rod and cone mutant mice and CBC polarity switch

Our data suggest that a rod-cone interaction induced polarity switches in ON and OFF CBCs and SACs (Fig. 2). We used rod or cone functional knockout mice to test the mechanisms of the polarity switch. We first used the *Gnat1^-/-^*mice in which rods are dysfunctional ^26, 27^. Recordings in these mice revealed that ON and OFF CBCs did not respond to light in the mesopic conditions (Fig. 3). In the photopic conditions, ON CBCs depolarized to light, and OFF CBCs hyperpolarized to light onset, exhibiting no signs of a polarity switch (Fig. 3a). These results suggest that rod photoreceptors induced the polarity switch.

**Fig. 3.**
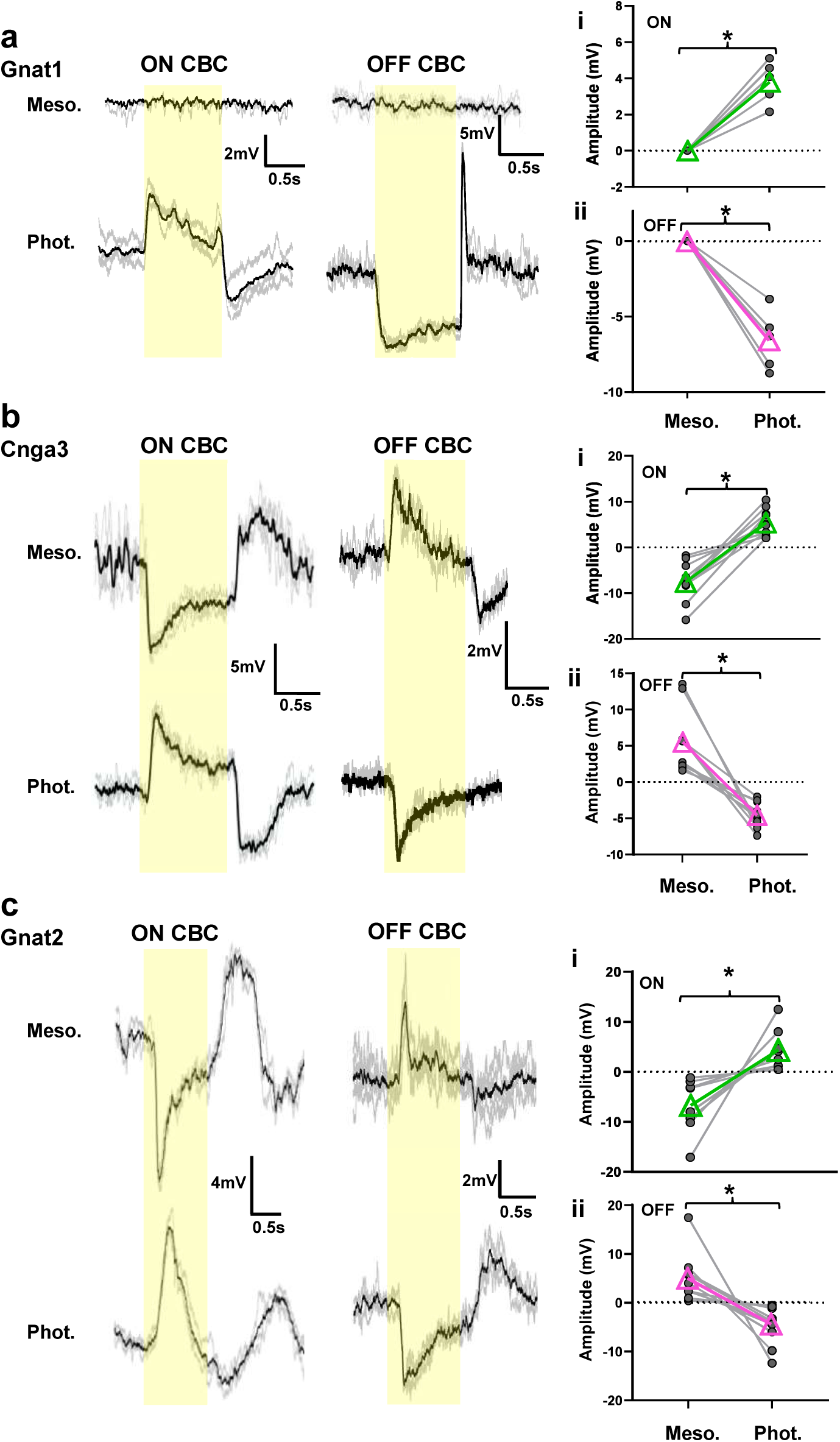
Rod and cone dysfunctional mutant mice and polarity switch. **a.** (*left*) L-EPSPs were recorded from an ON CBC in response to 1 log step stimulus (yellow bar) in *Gnat1^−/–^* at mesopic and photopic background luminances. This cell showed no response at mesopic and depolarized at the photopic background level, exhibiting no polarity switch. (*right*) L-EPSPs were recorded from an OFF CBC, showed no response at mesopic and hyperpolarized at the photopic light level, showing no polarity switch. A summary graph for peak amplitudes of L-EPSPs for ON CBCs (**i**) and OFF CBCs (**ii**) showed no response in mesopic but responded with correct sign in photopic. Here and subsequent panels, black circles indicate peak amplitudes in individual CBCs, and green (ON CBCs, n=10, p<0.05) and magenta (OFF CBCs, n=5, p<0.05) triangles show average responses. **b.** Representative L-EPSPs in ON and OFF CBCs in *Cnga3^-/-^* at mesopic and photopic conditions in response to 1 log step stimulus (yellow bar). Polarity switch was observed in mesopic and correct sign L-EPSPs were evoked in photopic conditions. The black trace is the average of the gray trials for all examples. **i.** A graph showing peak amplitudes for 13 ON CBCs at mesopic and photopic conditions (n=8, p<0.005). **ii**. OFF CBCs for same conditions as (**i**) (n=13, p<0.005). **c.** Representative L-EPSPs for an ON and OFF CBCs in *Gnat2*^−/–^ cone dysfunctional mice. **i.** Peak amplitudes for ON CBCs, which showed a polarity switch in mesopic and correct sign in photopic (black circles) (n=8, p<0.005). **ii.** Peak amplitudes for OFF CBCs for same conditions as (**i**) (n=10, p<0.005). All statistical tests in this figure are Wilcoxon *t*-test. Asterisks represents statistically significant effects.

Next, we used the *Cnga3^-/-^* mice in which cones are dysfunctional, but rods are intact ^28, 29^. In mesopic conditions, ON CBCs hyperpolarized at light onset, showing a polarity switch (Fig. 3b, Table 1), additional evidence that rods play a critical role in the polarity switch (Fig. 3a). Then, we tested the photopic condition. Because these mice did not possess functional cones, we expected either the polarity switch to remain, or not to respond to light due to rod bleaching.

However, in the photopic conditions, the same ON CBCs depolarized, which is the correct sign for those cells. The polarity switching response occurred again when the light level was decreased to mesopic conditions after the tissue was exposed to bright light conditions (Extended Data Fig. 3). Correspondingly, OFF CBCs depolarized in the mesopic and hyperpolarized in the photopic conditions (Fig. 3b). More than half of the ON and OFF CBCs we recorded exhibited a polarity switch (Table 1, *Cnga3^-/-^*).

We wondered whether the unexpected photopic response in *Cnga3^-/-^* mice occurred due to any strain conditions other than cone functional-knockout. We used another cone-dysfunctional mouse strain, *Gnat2^-/-^* mice ^30^ to examine if the unexpected photopic response still occurs in CBCs. In the mesopic condition, ON CBCs hyperpolarized and OFF CBCs depolarized, exhibiting a polarity switch, while their photopic responses displayed correct signs (Fig. 3c & Table 1, *Gnat2*^-/-^). These results confirmed that rods generate both the polarity switch and the correct sign signals.

The sign changes occurred within a few minutes after being adapted to a new ambient light level (Extended Data Fig. 3). A short adaptation was required from mesopic to photopic, and photopic to mesopic when repeated (Extended Data Fig. 3). Furthermore, the short adaptation was required for ON and OFF CBCs in wildtype, *Cnga3^-/-^*, and *Gnat2^-/-^* mice. These results suggest that the polarity switch and correct sign signals originated in rods but are evoked by different light levels and mediated through different mechanisms.

### Gap junctions mediate the correct sign but not the polarity switch

Rod-cone interactions might occur through rod-cone coupling ^31^. We used a gap junction blocker, MFA (50 µM or 100 µM), to examine whether the rod-cone coupling mediated the polarity switch signals. An OFF CBC exhibited a polarity switch in the mesopic and a correct sign in the photopic conditions (Fig. 4a). We applied MFA to observe light responses over 35 minutes in both light conditions. After 5 minutes, the polarity switch in the mesopic conditions did not change, but the hyperpolarizing response (correct sign) in bright light was reduced. Over 35 minutes of recording, the polarity switch in the mesopic condition stayed the same, while the hyperpolarization (correct sign) in the photopic condition changed to depolarization, a polarity switch (Fig. 4a). A similar trend was observed in many ON and OFF CBCs (Fig. 4b). These results suggest the following: (1) Polarity switch and correct sign signals are mediated to CBCs through distinct pathways. (2) Gap junctions, including rod-cone coupling and AII amacrine-ON CBC coupling, mediate correct sign signals but not the polarity switch signals. (3) Horizontal cells may contribute to generating correct signs in CBCs, whose signals spread through gap junctions among these cells ^32, 33^. We looked further into the contributions of horizontal cells to the polarity switch.

**Fig. 4.**
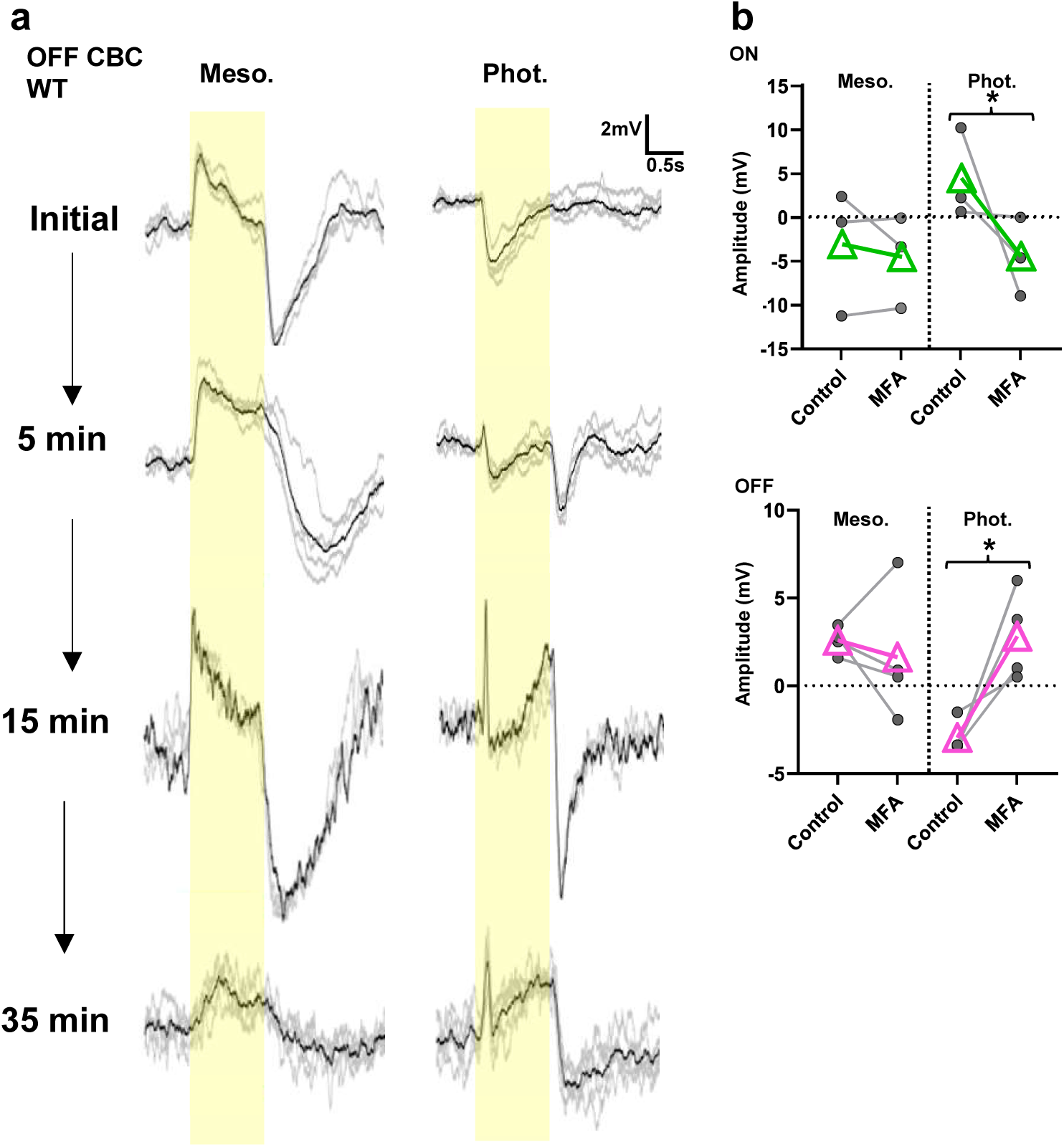
Gap junctions mediated correct sign but not polarity switch in ON and OFF CBCs. **a.** L-EPSPs in an OFF CBC in WT retina in response to 1 log unit step light (yellow bar) at mesopic background luminance before and during application of 100 µM MFA to block gap junctions. The black trace is the average of the gray individual traces for the condition. (*left*) L-EPSPs exhibited polarity switch in control, which was not changed as the drug took effect. (*right*) L-EPSPs in photopic background luminance. The hyperpolarizing L-EPSP was converted to depolarization as the drug took effect. **b.** (*upper*) A summary graph shows L-EPSP amplitudes from ON CBCs in two different background levels and before and after the MFA application (n=3, p<0.05). One ON cell did not show polarity switch initially, but the response was flipped after MFA. (*lower*) OFF CBCs showed similar results to the upper panel (n=4, p<0.05). All statistical tests in this figure are student t-test. Asterisks represents statistically significant effects.

### Horizontal cells and mGluR6 contributions to the polarity switch

Only one type of horizontal cell is reported in the mouse retina, which contacts cones with dendritic arbors and rods with their axon terminals ^34^. Cellular mechanisms of feedback (from horizontal to photoreceptors) and feedforward (horizontal to bipolar cells) signal transmission are not fully elucidated ^35–39^. Furthermore, positive and negative feedback occurs locally and globally from horizontal cells ^40^, suggesting that complicated modulatory effects occur in the OPL.

Nevertheless, we manipulated the horizontal cell activity by blocking photoreceptor synaptic input with an AMPA/kainate glutamate receptor antagonist, CNQX (15 µM) ^41, 42^. We tested this in *Gnat2^-/-^* mice in which CBC signals originated solely from rods. In these mice, OFF CBCs exhibited a polarity switch at the mesopic and a correct sign in the photopic conditions (Fig. 5a). The application of CNQX did not change the polarity switch in the mesopic. In contrast, the correct sign in photopic conditions was converted to a polarity switch by CNQX, depolarizing to light (Fig. 5a, n=4). Some types of OFF CBCs express glutamate receptors sensitive to CNQX ^9^, and CNQX abolished EPSPs of both polarity switch and correct signs (n=4, data not shown).

**Fig. 5.**
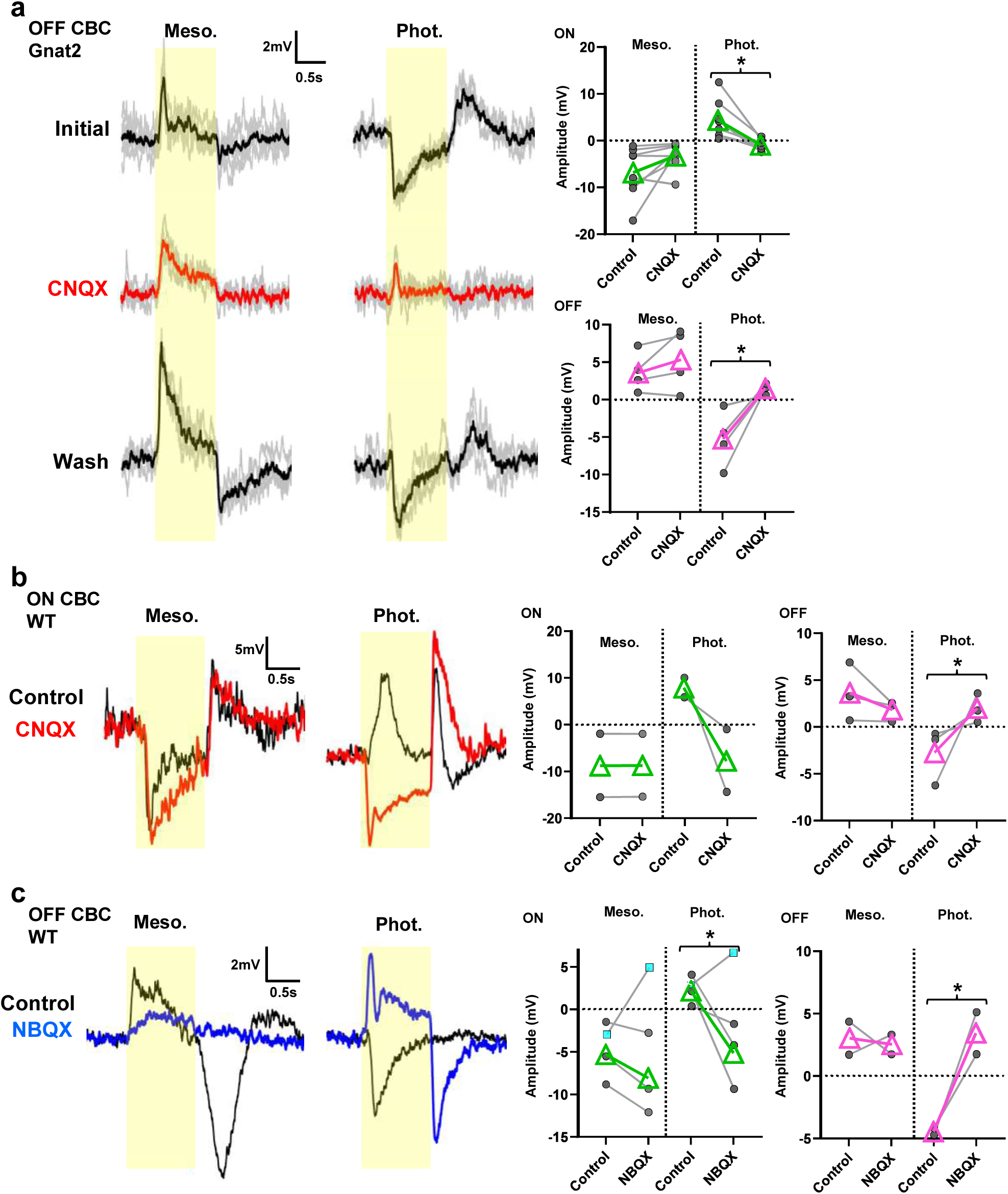
Blocking transmission from photoreceptors to horizontal cells reduced the correct sign but did not change polarity switch. **a.** L-EPSPs were recorded from an OFF CBC in *Gnat2*^−/–^ cone KO retina in response to 1 log unit step light for 1 second (yellow bar) in mesopic and photopic conditions and before, during, and after application of 15 μM CNQX, an AMPA/kainite receptor antagonist. While CNQX did not change flipped L-EPSP in mesopic, it converted the correct sign response in photopic to a polarity switch. (*Upper*) A graph shows L-EPSP amplitudes in 7 ON CBCs (Photopic, p<0.05), and lower graph shows amplitudes from 4 OFF CBCs (Photopic, p<0.05). **b.** L-EPSPs from an ON CBC from WT retina with the same stimulus in mesopic and photopic. CNQX did not change the polarity switch but converted correct sign to polarity switch in photopic conditions. Graphs summarize L-EPSP amplitudes from 2 ON and 3 OFF CBCs, showing a similar effect of CNQX. (OFF photopic, p<0.05) **c.** Similar effects were seen when using 10 μM NBQX, a potent AMPA receptor antagonist, in place of CNQX. L-EPSPs from an OFF CBC in WT retina are shown before and after application of NBQX. The polarity switch response (depolarization) at mesopic was not converted by NBQX, but correct sign signal (hyperpolarization) at photopic was flipped to depolarizing after NBQX application. 4 ON cells (3 cells had the polarity switch at the photopic level “black circles”, and 1 cell had the polarity switch at the mesopic level “cyan square”) and 2 OFF cells showed polarity switch at the photopic level (ON, n=3, p<0.05; OFF, n=2, p<0.05). All statistical tests in this figure are student t-test. Asterisks represents statistically significant effects.

Similarly, for ON CBCs with a polarity switch in mesopic conditions, CNQX switched the correct sign in photopic conditions without affecting the polarity switch in mesopic conditions (Fig. 5a). A similar CNQX effect was observed in WT mice, which possess both rods and cones; no changes for the polarity switch in mesopic, but CNQX converted the correct sign in photopic conditions to a polarity switch (Fig. 5b).

Furthermore, we examined an additional antagonist, NBQX (10 µM), a potent AMPA receptor antagonist, which was used for the photoreceptor-horizontal cell transmission blockade ^22^. The effect of NBQX was similar to the CNQX outcome: it had no effect on polarity switched signals in mesopic conditions but changed the correct sign to a polarity switch in photopic conditions (Fig. 5c). These results indicate that horizontal cells mediated the correct sign signals but did not mediate the polarity switch.

We then tested whether the polarity switching signals were mediated through glutamate receptors on bipolar cell dendrites. We used L-AP4 to block mGluR6-signaling in ON CBCs. L-AP4 abolished both EPSPs with a polarity switch and correct signs, while both signs of signals in OFF CBCs were unaffected by L-AP4 (Extended Data Fig. 4). The results are consistent with the L-AP4 application to SACs (Fig. 1h), and confirmed that both polarity switch and non-switch signals were mediated via glutamate receptors on CBC dendrites.

### Excitatory amino acid transporter 5 (EAAT5) was involved in generating the polarity switch

Rod and cone axon terminals express glutamate transporters, EAAT2 and EAAT5, to uptake glutamate released at the terminals ^43, 44^. EAAT5 has a large chloride conductance in conjunction with glutamate uptake, which could generate the inhibitory signals ^2, 45–47^. We used a glutamate transporter inhibitor, DL-TBOA (50 μM) to test whether EAAT5 generated the polarity switch. ON and OFF CBCs exhibited different signs of L-EPSPs in mesopic and photopic conditions in wildtype mice (Fig. 6). DL-TBOA corrected the polarity switch without affecting the correct sign signals (Fig. 6a,b). During the DL-TBOA application, the CBC membrane potential was not changed (n=14 CBCs, control: −42.6 ± 1.4 mV, TBOA: −42.4 ± 1.5 mV). These results suggest that EAAT5 and associated chloride conductance in cone terminals generate depolarizing light responses and induce the polarity switching signals in second-order neurons (Fig. 6c).

**Fig. 6.**
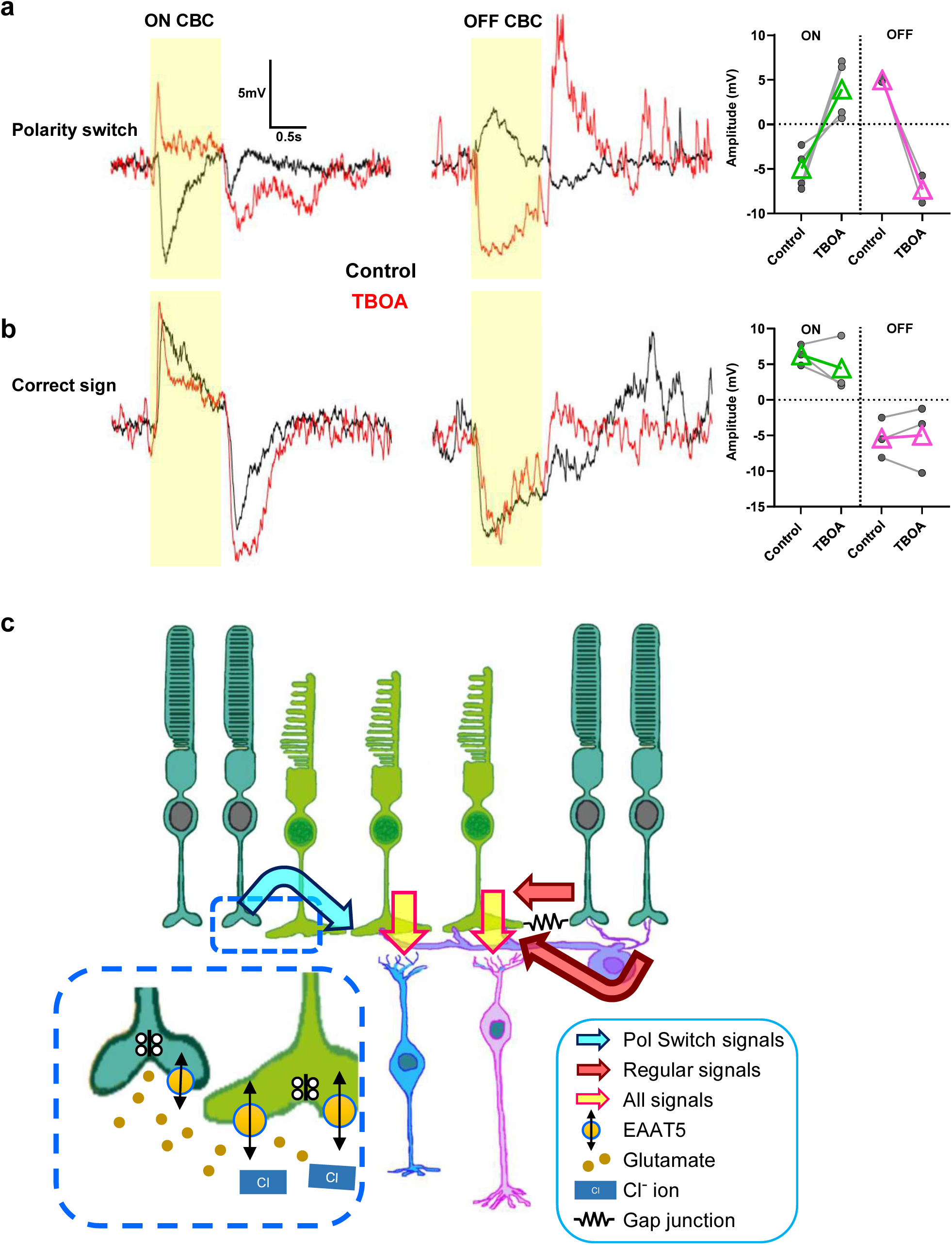
The EAATs mediated the polarity switch in CBCs, and a schematic model showing the polarity dichotomy pathways. **a.** An ON CBC and an OFF CBC exhibited L-EPSPs with polarity switch. DL-TBOA (50 μM) removed the polarity switch for 4 ON CBCs and 2 OFF CBC. **b.** L-EPSPs without polarity switch in an ON CBC and an OFF CBC was not changed by DL-TBOA. **c.** Graphical schematic of a working model for the polarity dichotomy pathways. (1) The correct sign L-EPSPs (red arrow) originate in rods and cones. Rod-originated signals are mediated through rod-cone coupling and horizontal cells to reach cones. Both rod- and cone-originated signals are then transmitted to ON and OFF CBCs (yellow arrows). (2) The polarity switch signals also originate in rods (blue arrow). Continuously released glutamate activates EAAT5 and Cl-currents in cones in dim light conditions, inhibiting cones. When light stimulus reduces glutamate release from rods, cone inhibition stops, depolarizing the cone terminals. This generates the L-EPSPs with polarity switches in ON and OFF CBCs. Cones are green, rods are dark cyan, horizontal cells are purple, an ON CBC is pink, and an OFF CBC is blue.

## Discussion

ON and OFF signals are generated at the first synapse of the retina between photoreceptors and bipolar cells. In the dark, photoreceptors are depolarized and continuously release glutamate. OFF bipolar cells receive glutamatergic inputs through AMPA and kainate glutamate receptors ^9, 48, 49^, and are depolarized in the dark. In contrast, ON bipolar cells express mGluR6 linking to TrpM1 channels ^50–52^, and are hyperpolarized in the dark but depolarized in the light as photoreceptor glutamate release is reduced. The ON and OFF dichotomy is separately transferred to ON and OFF sublaminae in the IPL, mediating the signals to ON and OFF amacrine and ganglion cells. This morphological and physiological relation is acknowledged as one of the fundamental rules in retinal neurophysiology ^5–7, 11, 12, 53, 54^, throughout the visual system ^55^, and over the ranges of light conditions from scotopic to photopic ^56, 57^. Despite the long-established consensus, we found that ON-OFF polarity switch occurs in retinal inner neurons.

There are only a few cases to date where this rule has been violated. Vlasits, et al. ^15^ reported that a polarity switch occurs in ON and OFF SACs. This switch even causes directional preference changes in direction-selective ganglion cells (DSGCs) ^58, 59^. Furthermore, Szikra, et al. ^22^ reported that rod photoreceptors exhibit polarity switching signals at high light, which is mediated by cone signaling through horizontal cells. For both cases, antagonistic surround through amacrine or horizontal cells was part of the underlying mechanisms. However, we used a relatively small spot of light to evoke light responses, and antagonistic surround was not the reason for the polarity switch (Figs. 1,5 and Extended Data Fig. 1).

A robust polarity switch occurred both in ON and OFF SACs as well as ON and OFF CBCs (Fig. 2). Polarity switch signals were evoked in mesopic, and correct sign signals were in photopic conditions, suggesting that rod-cone interaction induced the polarity switch. Because the sign switching occurred within a few minutes (Extended Data Fig. 3) and the pharmacological applications differentially blocked L-EPSPs (Figs. 4-6), we showed that two types of signals with opposite polarities were mediated through different mechanisms (Fig. 6c). Although the mechanisms are different, rods generate both polarities of signals (Fig. 3), and glutamate receptors on bipolar cell dendrites mediated both polarities (Extended Data Fig. 4).

L-EPSPs with correct sign were generated both by cones and rods. L-EPSPs generated by cones were transferred to CBCs and SACs, shown in photopic conditions, and in the rod-dysfunctional (*Gnat1-KO*) mice (Figs. 2-3). Rods also generated L-EPSPs with correct sign in CBCs, mediated through rod-signaling pathways and horizontal cells (Figs. 3-5). Three rod-signaling pathways have been reported, in which the primary and secondary pathways mediate rod signaling to ON and OFF CBCs through AII amacrine-ON CBC and rod-cone coupling, respectively ^60^. Our data with MFA revealed that the correct sign signals were mediated through gap junctions, corroborating rod-signaling pathways (Fig. 4).

Mutant mouse experiments revealed that rods generate the polarity switch signals (Fig. 3), which reach the CBCs through glutamate receptors on their dendrites (Fig. 1h and Extended Data Fig. 3). However, against our expectations, horizontal cells did not mediate the polarity switch (Extended Data Fig. 1 & Fig. 5), nor did gap junctions, including rod-cone coupling (Fig. 4). There is a direct synapse between rods and OFF CBCs, but not with ON CBCs ^60, 61^, which cannot explain the polarity switch in ON CBCs. Without any other putative mechanisms between rods and CBCs to mediate polarity switch signals, we decided to test whether glutamate transporters in photoreceptor terminals contributed to the signals. We found that application of the EAAT transporter blocker DL-TBOA removed the polarity switch without affecting correct sign EPSPs (Fig. 6).

Glutamate transporters, EAAT2 and EAAT5, are expressed by axon terminals of rods and cones ^43, 62^. Among those, EAAT5 has a large chloride conductance ^47^, which might affect the photoreceptor membrane potential and glutamate release. A potential explanation for the results with DL-TBOA is as follows: in dim-light conditions, glutamate is continuously released from rods and cones, inducing the glutamate spillover from rods to cones. This activates EAAT5 and the chloride current in cone terminals, inhibiting cones. When a light stimulus reduces the glutamate release from rods, EAAT5-linked chloride conductance is reduced, disinhibiting cones. This results in light-evoked cone depolarization and evoking the polarity switch signal in CBCs.

There are two critical points for this scenario to occur: (1) the spillover occurs among rods and cones and (2) the chloride conductance induces inhibitory currents in cones. Although a massive EAAT5 exists at the rod terminals to clean up released glutamate ^45^, EAAT5 saturates quickly, and spillover occurs, at least among cones ^63, 64^. Rod-to-cone glutamate spillover may also occur. For the second point, previous studies have measured the resting membrane potentials (RMP) as well as chloride reversal potential (E_Cl_). The RMP in cones in the mouse retina is between −45 and −55 mV ^65^, and the E_Cl_ has been reported in various species with a range of values: −42 to −55 mV ^66^. These potentials are close to each other, which may not induce any currents. However, photoreceptors are depolarized in dim conditions and may cause hyperpolarization upon EAAT5-chloride conductance activation.

DL-TBOA-sensitive EAATs on BC dendrites have been reported in the fish ^67^ and the mouse ^68^ retinas. If these EAATs generate chloride currents, they are potentially involved in producing a polarity switch. However, DL-TBOA did not change the membrane potentials in our recordings (Results section). Furthermore, the polarity switch occurred differently in ON and OFF CBCs (Fig. 2), which cannot be induced by EAAT-induced chloride currents if existed in postsynaptic sites. For these reasons, DL-TBOA-sensitive EAATs were located in presynaptic as reported earlier (Fig. 6c) ^43, 62^.

In our recordings, the polarity switch occurred only in approximately half of CBCs and SACs (Fig. 1, Table 1). This occurred both in ON and OFF cells, but we did not see cell type-dependency, nor retinal topographic distributions. We consider this might be due to the close proximity of the RMP and E_Cl_, resulting in a range of EAAT5-Cl-channel effects on cone membrane potentials across the retina. Our recordings revealed that the CBC polarity switch occurred more frequently in cone-KO than in WT mice (Table 1: Cnga3: 77%, Gnat2: 86%, WT: 52%). Cones in cone-KO mice do not change the RMP as the ambient light level changes. In contrast, cones in WT mice are hyperpolarized in bright conditions, diminishing the EAAT5-induced hyperpolarization and reducing the occurrence of CBC polarity switch.

Unlike previous studies with slice recordings ^8, 9^, here we found that more than half of CBCs exhibited a polarity switch (Table 1). We consider that the reason for the difference is the preparation: slice (previous) and wholemount (current) preparations. Wholemount preparations are more physiological than sliced preparations, which contain intact vertical and lateral neural connections. Interestingly, they preserve not only large neurons, such as horizontal cells, but also rod signaling. Rod photoreceptors saturate their responses at the high mesopic conditions: ∼10^4^ photons/µm^2^/s ^69, 70^. However, a couple of articles which used wholemount retinal preparations, claim that rods do not saturate ^71–73^. In our previous study with slice preparations, we did not observe any light responses in rod bipolar cells when we applied the background light at the “rod-saturating” light level ^8^. In contrast, rod-activated responses are evoked from mesopic to photopic conditions in *Cnga3^-/-^* and *Gnat2^-/-^* mice (Fig. 6), and rod bipolar cells exhibited light responses in both mesopic and photopic conditions (Extended Data Fig. 4). There might be chemical mechanisms to support rod signaling from the response saturation in *ex vivo* preparations. Presumably, well-preserved rods generate a polarity switch to provide impact on cone-signaling in mesopic to even in the photopic ranges.

The polarity switch is potentially critical to suppress signals in CBCs and SACs in dim light conditions when rod-signaling is dominant. It may also increase L-EPSPs during the transition from mesopic to photopic conditions by increasing the driving force. Furthermore, instantaneous polarity switching with short adaptation may be critical for the saccadic shift or continuously moving natural scene imaging. Future research will require clarifying the significance of the ON and OFF polarity switches in retinal neurons.

## Methods

### Ethical approval

All conducted animal studies adhered to ethical protocols approved by their Institutional Animal Care and Use Committee at Wayne State University (protocol no. 23-11-6310). Experiments were in line with the ARVO Statement for the Use of Animals in Ophthalmic and Visual Research. All procedures were taken to minimize animal suffering and maintain the physiological conditions of the retinal tissues.

### Retinal preparation

The experimental techniques used were described in detail in our previous studies ^8, 24^. Briefly, both male and female mice, aged 4 – 12 weeks, were used including the C57BL/6J strain (RRID: IMSR_JAX:000664), Ai9 (RCL-tdT) (RRID: IMSR_JAX:007909), R26R-EYFP (RRID: IMSR_JAX: 006148); Jackson Laboratory, Bar Harbor, ME, USA. In addition, rod knockout mice (*Gnat1^-/-^* ^74^) and cone knockout mice (*Cnga3^-/-^*) both gifted by Dr. Samar Hattar, and cone knockout mice (*Gnat2^−/–^* ^30^ gifted by Dr. Marie Burns) were also used. All mice were dark adapted overnight before being euthanized using carbon dioxide followed by cervical dislocation and eye enucleation. Retinal tissues were isolated under a stereo microscope. All steps were conducted in dark-adapted conditions utilizing infrared viewers. The dissecting solution was maintained at a cool temperature and continuously oxygenated. The dissections were performed in a HEPES-buffered medium (see solutions section for more details), and the retinal tissues were kept in an oxygenated dark box at room temperature.

### Whole-cell recordings

We performed whole-cell patch clamp recordings from bipolar cells and SAC somas in wholemount retinal preparations using an upright microscope (Slicescope Pro 2000, Scientifica, UK) with a CCD camera (Retiga-2000R, Q-Imaging, Canada). The tissues were stabilized with a platinum horseshoe net and nylon wires. Light-evoked postsynaptic potentials (L-EPSPs) were recorded at the resting membrane potential. Recordings were conducted at temperatures between 32-34°C, and liquid junction potentials (∼7.4mV) were adjusted post-recording.

Electrodes were made from borosilicate glass (1B150F-4; WPI, FL, USA) using a P1000 Puller (Sutter Instruments, CA, USA), with resistances ranging from 6 to 12 MΩ. Clampex and MultiClamp 700B (Molecular Devices, CA, USA) were utilized to generate waveforms, acquire data, and control an LED light stimuli (Cool LED, UK). Data was digitized with an Axon Digidata 1550B (Molecular Devices), filtered at 1 kHz using the four-pole Bessel filter on the MultiClamp 700B, and sampled at rates between 2 and 5 kHz.

### Solutions and Drugs

Retinal dissections were carried out in a HEPES-buffered extracellular solution containing 115 mM NaCl, 2.5 mM KCl, 2.5 mM CaCl_2_, 1.0 mM MgCl_2_, 10 mM HEPES, and 28 mM glucose, adjusted to a pH of 7.37 using NaOH. Physiological recordings were conducted in Ames’ medium, which was buffered with NaHCO_3_ (Millipore-Sigma, St. Louis, MO, USA) and aerated with 95% O2 and 5% CO_2_, maintaining a pH of 7.4 at a temperature range of 30-33°C. The intracellular solution used included 110 mM KMeSO_3_, 1.0 mM MgCl_2_, 10.0 mM HEPES, 4.0 mM EGTA, 5.0 mM NaCl, 5.0 mM KCl, 4 mM ATP-Mg, and 0.5 mM GTP-Mg, adjusted to a pH of 7.2 using KOH. We substituted KMeSO_3_ with CsMeSO_3_ for the recording in voltage-clamp mode.

The following pharmacological agents were bath applied: a glycine receptor antagonist, strychnine (1 μM, Sigma, St. Louis, MO), a GABA_A_ receptor antagonist, bicuculline methobromide (50 μM, Enzo Life Sciences, Farmingdale, NY), a GABA_B_ receptor antagonist CGP-52432 (10 μM, Tocris, Minneapolis, MN), and a GABA_C_ receptor antagonist, (1,2,5,6-tetrahydropyridin-4-yl) methylphosphinic acid hydrate (TPMPA, 50 μM, Tocris). L-(+)-2-Amino-4-phosphonobutyric acid (L-AP4, 10 μM, Tocris) was used as an mGluR6 agonist. To suppress Kv3 channel activity, 1 mM tetraethylammonium (TEA, Sigma) was bath applied or 10 mM TEA was included in intracellular solution ^13, 75^. 20 mM HEPES was applied to block horizontal cell feedback ^36, 76^ and 20 mM sucrose was added in Ames’ solution and 10 mM KCl was added to intracellular solution for control recordings. AMPA and kainite glutamate receptor antagonists, 6-cyano-7-nitroquinoxaline-2,3-dione (CNQX) (15 µM, Tocris) and 2,3-Dioxo-6-nitro-1,2,3,4-tetrahydrobenzo[f]quinoxaline-7-sulfonamide (NBQX) (10 μM, Tocris), were bath applied to block photoreceptor - horizontal cell transmission. To block gap junction, meclofenamic acid (MFA) (50-100 µM, Sigma) was used. Lastly, we used DL-threo-β-Benzyloxyaspartic acid (DL-TBOA, 50 μM, Tocris) to inhibit EAAT5.

### Light stimulation

A green (500 nm) spot light was directed through a 60× objective lens to illuminate photoreceptors around the recorded cells. We covered an area of 150 μm diameter, which is approximately the receptive field size for a starburst amacrine cell (SAC), and slightly larger than the receptive field center of a bipolar cell ^12, 77, 78^. We defined the mesopic light levels range as between 10^2^-10^4^ photons/ μm^2^/s and photopic range as greater than 10^5^ photons/ μm^2^/s. The preparations were adapted to background illumination at the mesopic range for approximately 10 minutes prior to recording. A step of light (with 20% or 85% Weber contrast, lasting 1 second) was projected on top of the background illumination. The light response at each background was checked twice for SAC and bipolar cells recordings: after incrementing backgrounds, we would return to the preceding lower intensity background to check the original response; one-minute intervals were used between backgrounds.

### Morphological Identification

A patch clamp pipette was filled with a fluorescent dye, sulforhodamine B (0.005%, Sigma) or Alexa 488 (0.003%, ThermoFisher Scientific, Waltham, MA). Immediately following electrophysiological recordings, images of bipolar cells were visualized and their morphological types confirmed by their dendritic characterizes (SAC) and axonal ramification patterns (BC) in the IPL ^25^. Images of both axon terminals and ChAT-bands were captured and recorded using a CCD camera.

For retinal tissues without ChAT-band fluorescence, such as wildtype and *Cnga3^-/-^*, we included Neurobiotin (0.5%, Vector Lab, CA, USA) in the pipette solution along with a fluorescent dye ^8, 9^. To visualize cells with Neurobiotin, the preparation was fixed with 4% paraformaldehyde for 30 minutes, then incubated overnight with streptavidin-conjugated Alexa 488 (1:200, Thermo Fisher Scientific) and an anti-choline acetyltransferase (ChAT) antibody (1:200, AB144P, Millipore, MA, USA). Subsequently, the preparation was incubated with the secondary antibody for 2 hours at room temperature. The preparation was then observed using a confocal microscope (TCS SP8, Leica, Germany). Bipolar cell types were identified based on previous descriptions ^8, 9, 79^.

### Data analysis and statistics

For step-pulse light-evoked L-EPSPs, we measured the peak amplitude (in millivolts) using Clampfit software (Molecular Devices). All values are presented as the mean ± SEM. SAC and bipolar cell responses to pharmacological agents was evaluated by Student *t*-test (paired) and the Repeated Measures ANOVA with Tukey’s multiple comparison. Bipolar cell polarity switch was evaluated using the Wilcoxon signed ranks test (pared, two-tail). Light levels and polarity switches were compared using the One-Way ANOVA with Tukey’s multiple comparison. All statistics were conducted using Prism 10.4 (GraphPad Software Inc., CA). The significance level was set at a threshold of p<0.05.

## Data Availability

The data supporting this Article are available from the corresponding author upon reasonable request. <u>Source data</u> are provided with this paper.

## Acknowledgements

We would like to thank Drs. Dao-Qi Zhang and Samar Hattar for providing photoreceptor KO mice. We also thank Dr. Manoranjan Santra for genotyping support and Mr. Bashir Khatib-Shahidi and Mr. Goichi Suganuma for technical assistance. We are grateful to generous research funding: NIH EY028915 (TI), NIH EY032917 (TI), NIH EY004068 (Vision Core), Rumble Fellowship (JB), and RPB grant.

## Author Contributions

C.B.H. and T.I. were responsible for conceptualizing the study. D.L.B., A.R.H., A.S. J.M.B. and C.B.H. performed patch clamp study and analyzed data. Y.U. and E.C.S. advised on light adaptations and provided photoreceptor mutant mouse analysis. T.I. wrote original manuscript. D.L.B., A.R.H., A.S., and C.B.H. generated figures and conducted statistical analysis. All authors reviewed and edited the manuscript.

## Materials & Correspondence

Material and correspondence requests should addressed to the corresponding author, Tomomi Ichinose, MD, PhD.

