## Supplemental figures for "Rod photoreceptors control the ON vs OFF polarity of cone-signaling neurons"

### Extended Data Figure 1

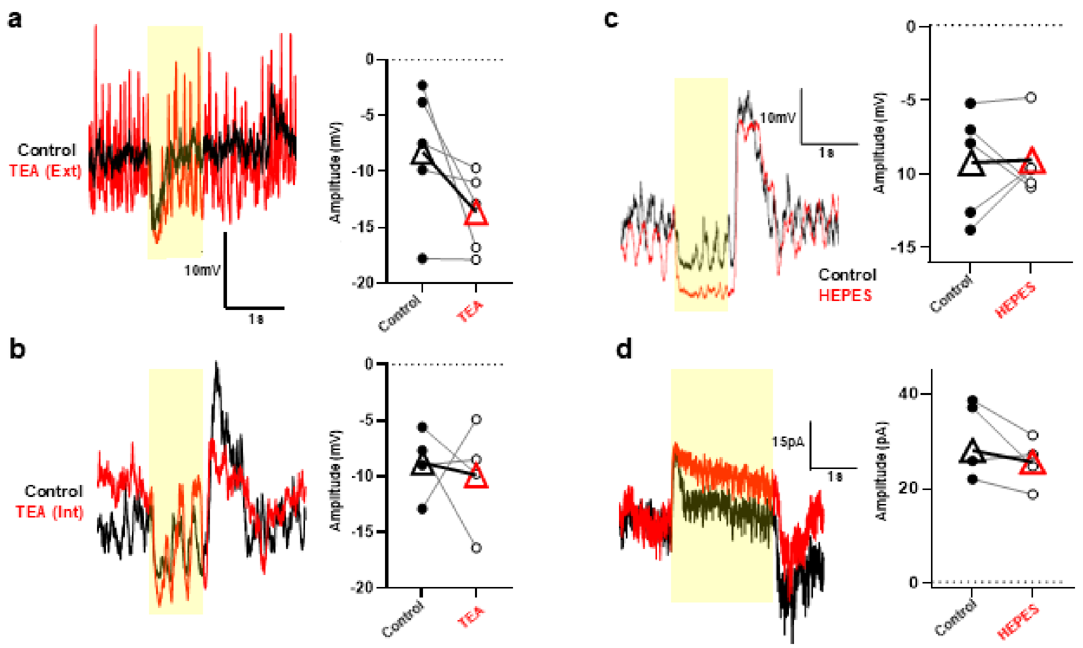


Extended Data Fig. 1

Kv3 channels and horizontal cells did not affect the polarity switch in ON SACs. **a.** A hyperpolarizing ON SAC before (black) and after (red) application of 1 mM TEA in the bath, blocking the Kv3 channels. A graph summarizes the L-EPSP amplitudes (n=5, p>0.05). **b.** A hyperpolarizing ON SAC with 10 mM TEA in the pipette solution. The black trace represents the response at break-in while the red trace represents the light response after 30 minutes. A graph summarizes the L-EPSP amplitudes (n=4, p>0.05). **c.** Representative averaged L-EPSPs to 1 second 20% contrast step light at mesopic conditions (yellow bar), showing ON SAC light responses before and after 20 mM HEPES. Polarity switch was not corrected by the HEPES applications. A summary graph showing the effects of HEPES on L-EPSPs. Black circles show individual ON SAC amplitudes, triangles show average responses. White circles indicate polarity flipping ON SACs after application of HEPES and red triangle is average of those cells only. **d.** Representative average voltage clamp traces to 2 second 20% contrast step light at mesopic, showing ON SAC light responses before and after 20 mM HEPES. The outward current in ON SACs were unchanged after the HEPES application. A summary graph showing effects of 20 mM HEPES with voltage clamp in 7 ON SACs. Black circles show individual amplitudes of ON SACs and black triangles are the average of those responses. All statistical tests in this figure are Wilcoxon *t*-test. Asterisks represent statistically significant effects.

Extended Data Figure 2

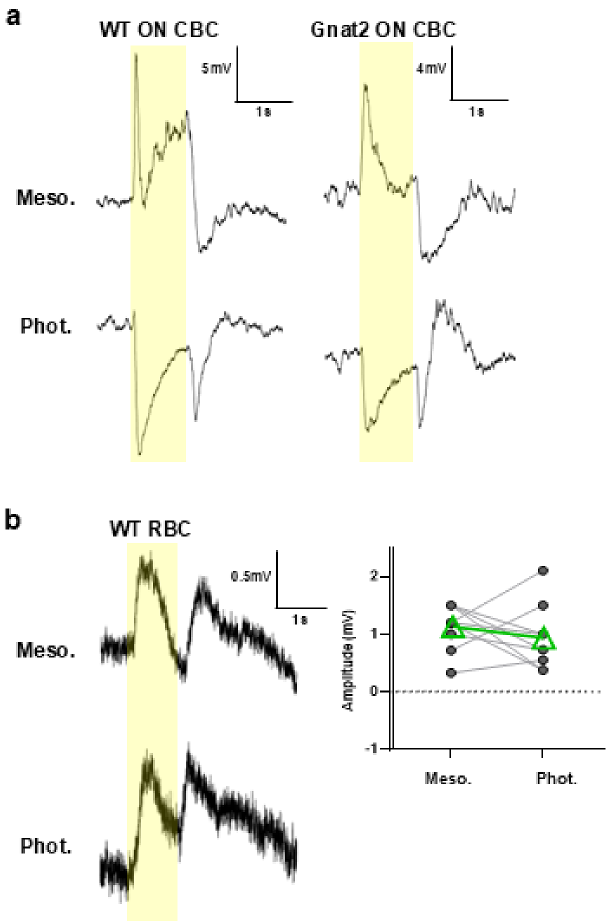


Extended Data Fig. 2

Some Cone bipolar cells (CBC) switch the opposite way and Rod Bipolar Cells (RBC) respond in photopic conditions without showing polarity switch. **a.** A representative L-EPSPs recording of 2 ON CBCs in different mouse models that underwent polarity switch in an opposite way, correct sign in mesopic and switched in photopic conditions. **b.** A representative L-EPSPs recording from an RBC in response to 1 log step stimulus (yellow bar) at mesopic (upper) and photopic (lower) background luminance. This cell depolarized in response to light onset to both light levels showing the expected sign and no polarity switch was noted.  A summary graph showing peak amplitudes of L-EPSPs of the RBCs in mesopic and photopic background luminance. RBCs did not show polarity switch (n=6, $p>0.05$). All statistical tests in this figure are Wilcoxon *t*-test. Asterisks represents statistically significant effects.

Extended Data Figure 3

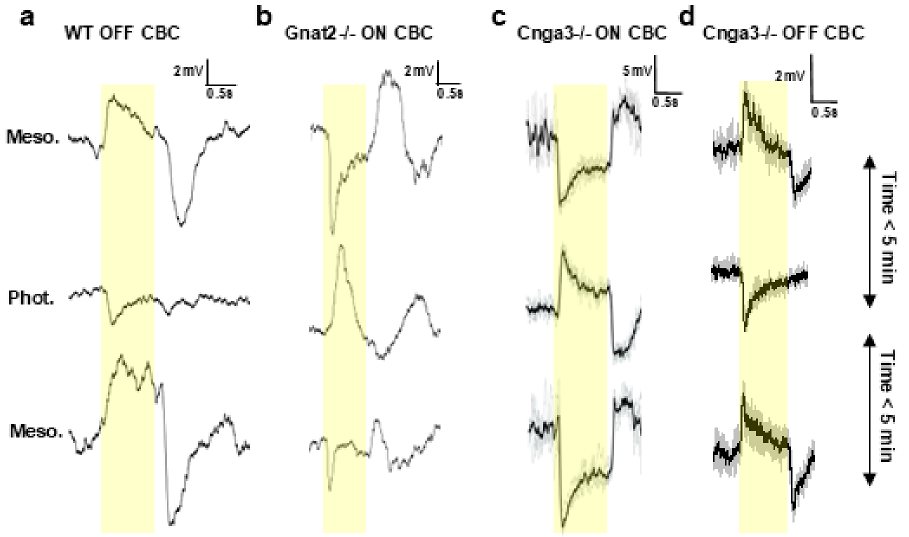


Extended Data Fig. 3

CBC polarity switch occurred within a few minutes. **a.** An OFF CBC initially depolarized in mesopic, switched the polarity in photopic, and switched back to depolarization in mesopic conditions. The polarity switch occurred within 5 minutes from the start of each condition to the next. **b.** Same conditions as **a.** for an ON CBC in *Gnat2*$^{-/-}$ cone KO retina showed similar fast changes in polarity. **c.** Same conditions as **a.** for an ON CBC in *Cnga3*$^{-/-}$ cone KO showed similar results. **d.** Same conditions as **a.** for an OFF CBC in *Cnga3*$^{-/-}$cone KO showed similar results.

Extended Data Figure 4

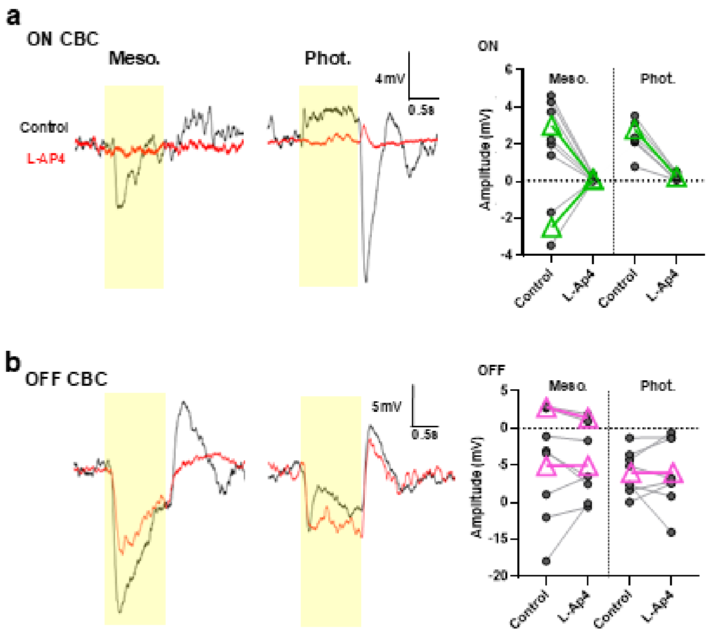


Extended Data Fig. 4

ON and OFF CBCs with L-AP4. **a.** A L-EPSPs from an ON CBC in WT retina in response to 1 log unit step light (yellow bar) at mesopic and photopic conditions before (black trace) and after application of 10 µM L-AP4 (red trace). L-EPSPs were entirely eliminated at both luminances. 2 ON CBCs showed initial polarity switches while 6 ON CBCs showed the correct sign to mesopic luminance level and both types showed the correct sign for photopic luminance level. All responses of ON CBCs were eliminated with L-AP4. Green triangles indicate average of individual responses. **b.** L-EPSPs from an OFF bipolar cell with same stimulus conditions as **a.** in the presence and absence of L-AP4. 2 OFF CBCs showed similar polarity switches while 6 cells showed the correct sign to mesopic luminance levels and both types showed the correct sign at photopic luminance levels. None of the responses were switched or eliminated by L-AP4. Magenta triangles indicate average of responses.
